# TWEAK Signaling Drives the Transition from Psoriasis to Atopic Dermatitis-like Inflammation in Paradoxical Skin Reactions

**DOI:** 10.1101/2025.06.19.660481

**Authors:** Sahiti Marella, Rachael Bogle, Haihan Zhang, Jennifer Fox, Joseph Kirma, Xianying Xing, Benjamin Klein, Vincent van Drongelen, Zoey Feister, Fadia Zagairy, Mira Hamed, Rundong Jiang, Poulami Dey, J. Michelle Kahlenberg, Allison C. Billi, Lam C. Tsoi, Johann E. Gudjonsson, Eran Cohen Barak

## Abstract

Targeted biologics have significantly advanced the treatment of inflammatory skin diseases such as psoriasis; however, some patients paradoxically develop eczematous skin reactions during or after anti-TNFα, IL-17, or IL-23 therapy. Although these paradoxical reactions resemble atopic dermatitis clinically and histologically, the molecular mechanisms that drive their development are not fully understood. Here, we generated high-resolution cellular and spatial maps of healthy skin, psoriasis, atopic dermatitis, and paradoxical reactions using single-cell RNA sequencing, spatial transcriptomics, immunohistochemistry, and in vitro assays. In paradoxical reactions, we identified a distinct transcriptional landscape characterized by myeloid and T-cell expansion and an altered keratinocyte phenotype shaped by TWEAK signaling. Mechanistically, we showed that TWEAK synergizes with IL-13 to drive the Th2/type I interferon-polarized epithelial program. Notably, anti-TNFα therapy induced TWEAK gene expression in myeloid cells, suggesting a compensatory inflammatory circuit. Together, these findings identify the TWEAK-IL-13 axis as a central driver of paradoxical skin reactions and provide a mechanistic framework for how cytokine blockade may rewire cutaneous immune responses.

One Sentence Summary

The TWEAK-IL-13 signaling axis is a key driver of immune reprogramming underlying paradoxical skin reactions.

## INTRODUCTION

Psoriasis (PP) is a chronic inflammatory skin disease characterized by epidermal hyperproliferation, abnormal keratinocyte differentiation, and robust immune cell infiltration, particularly in T and myeloid cells (1–3). It affects approximately 3% of the U.S. population and more than 125 million individuals worldwide, representing a significant burden on global health and quality of life (1, 4–6). Treatment strategies for PP range from topical steroids and systemic agents to biologics, which selectively target key inflammatory cytokines, such as tumor necrosis factor (TNFα), interleukin-17 (IL-17), and interleukin-23 (IL-23) (7, 8). These treatment strategies have significantly improved disease management in many patients, yet paradoxical reactions (PR) and new-onset or exacerbated inflammatory skin eruptions that resemble atopic dermatitis (AD) have emerged as perplexing and increasingly recognized adverse effect (9–13).

Unlike classical PP, which is primarily driven by the Th1 and Th17 cytokine pathways, AD is associated with a Th2-skewed immune response marked by elevated levels of IL-4, IL-5, and IL-13 (14–18). Some patients receiving TNFα/IL-17/IL-23 inhibitors for PP develop paradoxical eczematous skin eruptions that clinically and histologically resemble AD (19–21). Clinically, patients with suspected PR can present with diverse morphologies such as plaques, patches, xerosis, lichenification, and prurigo (22). Histopathologic features may include eosinophilia, spongiosis, and epidermal hyperplasia, while laboratory findings often show elevated serum immunoglobulin E (IgE) levels (22). Despite their clinical and histological resemblance to AD, PRs are uniquely marked by a robust type I interferon (IFN-I) gene signature, indicating innate immune activation, and supporting a distinct immunopathologic program (9, 14, 23).

The underlying molecular and transcriptional mechanisms driving PRs remain incompletely understood; however, one proposed mechanism is that pharmacological inhibition of Th1/Th17 pathways may relieve suppressive constraints on both Th2 and IFN-I pathways, leading to compensatory immune activation in susceptible individuals. We hypothesized that biologically induced immune rewiring may engage in novel ligand-receptor interactions between immune cells and keratinocytes. We sought to understand the spatial organization, cellular identities, and signaling pathways underlying this paradoxical skin inflammation.

## RESULTS

### Single-cell profiling of skin from NL, PP, PR, and AD skin reveals distinct cellular composition and histologic features in PR

To elucidate the cellular and molecular mechanisms underlying PR, we performed scRNA-seq, spatial transcriptomics, and histopathological analyses of skin biopsies from NL (n = 4), PP (n = 4), AD ( n = 4), and PR (n = 8) patients (Fig. 1A). Culprit medications included biologics for all classes (anti-TNFα = 2, Anti-IL-17 = 4, Anti-IL-23 = 2; Supplementary Table S1). Representative clinical images (n = 5 patients) highlight the distinct morphological features of PR skin, such as ill-defined erythematous plaques, pruritic skin-colored plaques covered with scales and crusts, and pruritic eczematous rash (Fig. 1B–F) (complete description of the PR study cohort is summarized in Table S1). Integration of single-cell data across disease groups identified 12 major skin cell populations: keratinocytes, fibroblasts, myeloid cells, T cells, melanocytes, mast cells, follicle cells, eccrine cells, adipocytes, pericytes, endothelial cells, and lymphatic endothelial cells (Fig. G-H). These cell types were defined by lineage markers such as *KRT14*, *KRT10* and *KRT1* (keratinocytes); *COL1A1*, *COL1A2*, and *COL3A1* (fibroblasts); *IL7R*, *PTPRC*, and *TRAC* (T cells); and *CD74*, *CLEC10A*, and *LYZ* (myeloid cells), and were consistently distributed across donors (Fig. 1I). Cell type composition analysis revealed marked shifts in specific compartments across disease states (Fig. 1H). While keratinocytes remained the dominant cell type in all groups, PR skin exhibited notable increases in T and myeloid cells relative to NL and PP samples, suggesting distinct immune activation pathways. Histopathological evaluation showed that PR was characterized by acanthosis, parakeratosis, varying degrees of spongiosis, and robust dermal immune infiltrates, further supporting the hybrid inflammatory phenotype unique to PR compared to NL, PP, and AD (Fig. 1J-M). Collectively, these data demonstrate that PR lesions exhibit a unique clinical presentation and distinct cellular and transcriptional landscape characterized by fibroinflammatory activation and augmented innate and adaptive immune responses, setting them apart from classical PP and AD.

**Fig 1.**
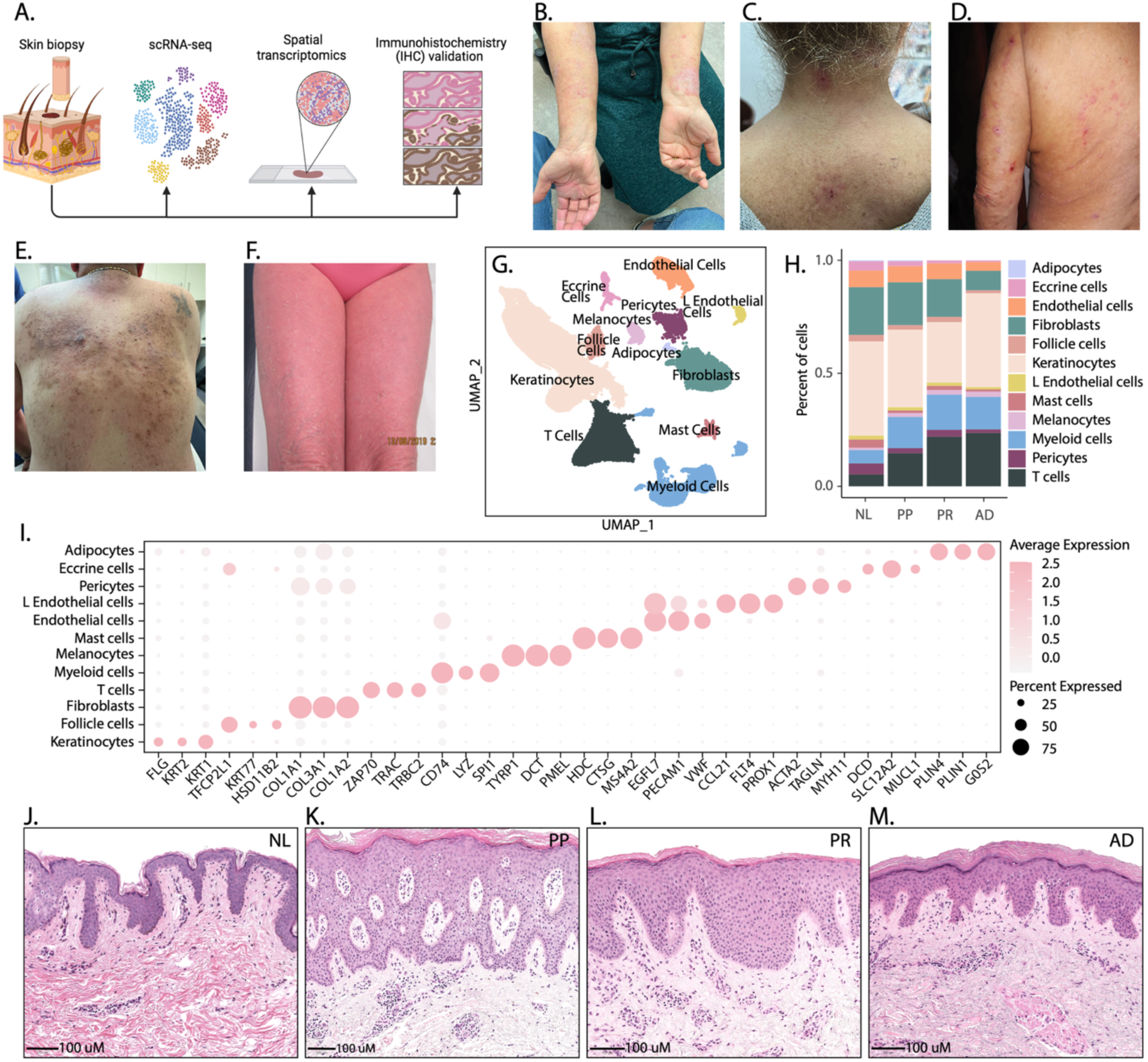
Single-cell profiling of skin from NL, PP, PR, and AD skin reveals distinct cellular composition and histologic features in PR. A. Summary diagram of study design representing skin biopsy collection, scRNA-seq, spatial transcriptomics, and immunohistochemistry (IHC) validation; (B-F) Clinical photos of patients with PR: B. Patient who developed PR following adalimumab treatment presented ill-defined erythematous plaques covered with scale on palms, soles, arms and lower limbs; C. Patient who presented with new onset of pruritic skin-colored plaques covered with scale and crusts following adalimumab treatment; D. Patient who presented with pruritic eczematous rash on trunk and extremities following ixekizumab treatment; E. Patient who presented with skin-colored papules coalescing to plaques covered with scale crust on the back following treatment with seckukinumab; F. Patient who presented with highly pruritic erythroderma following treatment with ixekizumab and PR persisted despite switching to gueselkumab; G. Uniform manifold approximation and projection (UMAP) visualization depicting integrated single-cell RNA sequencing (scRNA-seq) data from skin biopsies across disease groups; H. Cell composition analysis across disease groups showing the relative proportions (percentage) of each cell type across disease groups (NL, PP, PR, and AD); I. Dot plot visualization of average expression of top 3 key canonical cell type marker genes within major cell types; J-L. Representative brightfield hematoxylin & Eosin staining photomicrographs of skin biopsies of each disease group (NL, PP, PR, and AD).

### PR skin displays unique transcriptional and cell-signaling profiles, including enhanced IL- 4/IL-13 and TWEAK signaling

To identify transcriptional signatures specific to PR, we conducted pseudobulk differential expression analysis across skin biopsies from NL, PP, PR, and AD to identify the upregulated genes and pathways across groups. This analysis uncovered 674 genes that were uniquely upregulated and differentially expressed in PR compared with all other groups (Fig. 2A). Reactome pathway enrichment analysis on these PR-specific genes highlighted significant enrichment of immune-related pathways, prominently featuring the “Immune System” and underscoring the immunologic nature of paradoxical skin inflammation (Fig. 2B). Next, we performed deeper analysis of “Immune System” pathways across disease groups and identified enrichment of Interleukin-4 (IL-4) and Interleukin-13 (IL-13) signaling pathways in PR and AD (Fig. 2C), highlighting shared Th2-related immune mechanisms. Interestingly, PR skin uniquely demonstrated significant enrichment of the TWEAK (TNF-like weak inducer of apoptosis (TWEAK) signaling pathway (Fig. 2D), which has been implicated in several intracellular cascades, including activation of the nuclear factor-κB (NF-κB) pathway (24). TWEAK is a multifunctional proinflammatory cytokine that exerts its effects through its receptor, fibroblast growth factor-inducible 14 (FN14), to regulate diverse cellular processes such as proliferation, migration, apoptosis, and inflammation (24). Notably, emerging evidence has linked the TWEAK– FN14 signaling axis to the pathogenesis of both psoriasis and atopic dermatitis, highlighting its potential relevance in cutaneous inflammatory diseases. Given these associations, we sought to investigate the specific role of TWEAK signaling in driving the unique inflammatory milieu observed in PR. To identify the cell types that predominantly expressed the PR-specific transcriptional signature, we generated a gene module score from the 674 PR-specific DEGs and generated violin plots from the scRNA-seq data. We observed strong enrichment of the PR DEG module score within immune cells, most prominently in myeloid cells (Fig. 2E), indicating that these cells are key effectors in paradoxical inflammatory responses. Given the specific enrichment of the TWEAK signaling pathway in PR, we next assessed the cell type-specific expression of TWEAK (*TNFSF12*) and FN14 (*TNFRSF12A*). Feature plots of both *TNFSF12* (Fig. 2F) and *TNFRSF12A* (Fig. 2G) showed increased expression levels in PR skin, with robust expression in immune cell clusters compared to NL and PP, and partially overlapping with the patterns observed in AD. This heightened ligand–receptor co-expression further validates the active role of TWEAK signaling in PR. Finally, we characterized the cell-cell interaction landscape across disease groups using Circos plots to visualize intercellular communication networks (Fig. 2H-K). PR and AD, albeit to a lesser extent, display unique interaction profiles involving enhanced intracellular communication between myeloid cells, T cells, and stromal populations.

**Fig 2.**
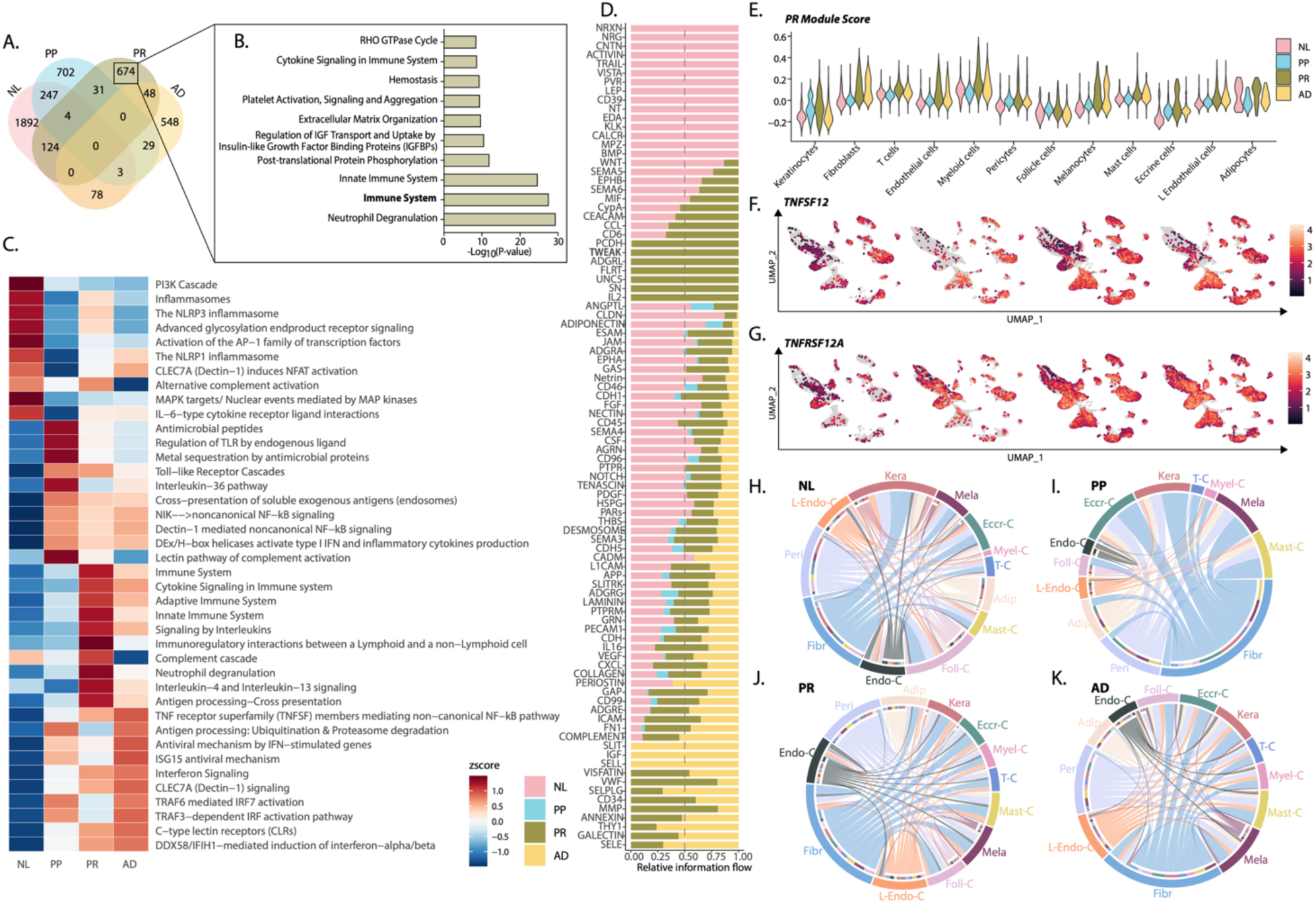
PR skin displays unique transcriptional and cell-signaling profiles, including enhanced IL-4/IL-13 and TWEAK signaling. A. Venn diagram illustrating common and unique upregulated pseudobulk-derived differentially expressed genes (DEGs) identified across disease groups (NL, PP, PR, and AD; B. Bar plot depicting the top 10 Reactome pathways enriched among the 674 PR-specific DEGs, highlighting “Immune System’ pathway; C. Top enriched “Immune System”-associated Reactome pathways across disease groups (NL, PP, PR, and AD); D. Top CellChat pathways enriched across disease groups, highlighting “TWEAK” signaling as uniquely enriched in PR; E. Violin plots across cell types and disease groups showing module scores generated from 674 PR-specific DEGs; (F-G) scRNA-seq feature plots across disease groups showing average expression: F. *TNFSF12* feature plot; G. *TNFRSF12A* feature plot; H-K. Circos plots visualizing predicted cell-cell communication networks across disease groups (NL, PP, PR, and AD).

### Myeloid–keratinocyte interactions drive paradoxical inflammatory responses via TWEAK signaling and enhanced cytokine communication

Given the significant enrichment of myeloid-driven immune pathways observed in PR, we investigated cell-specific interactions between myeloid cells and keratinocytes across disease conditions using Circos visualization. PR skin exhibited increased keratinocyte–myeloid interactions compared with NL, PP, and AD skin (Fig. 3A). These altered communication patterns suggest that myeloid-derived signals play pivotal roles in regulating keratinocyte activation and epidermal inflammation in PR. To validate these findings, we employed spatial transcriptomics to assess the localization of key inflammatory mediators in skin biopsies across disease groups. We first identified the major cell types, including keratinocytes, melanocytes, myeloid cells, pericytes, fibroblasts, endothelial cells, and T cells, across the disease groups (Figure 3B). Next, we performed gene expression analysis to assess the spatial distribution of *TNFSF12* and its *TNFRSF12A*, along with cytokines including *IL-4, IL-13, IL-17A, IL-17F, TNF, IFNG, IL-6, CCL22*, and *CCL17* in the PR skin (Fig. 3C-F). As anticipated, *IL-17A* and *IL-17F*, hallmark Th17 cytokines, were elevated in PP, whereas *IL-4* and *IL-13*, canonical Th2 cytokines, were upregulated in AD. Notably, *TNFRSF12A* was predominantly expressed by keratinocytes, whereas *TNFSF12* was enriched in myeloid cells, further highlighting the potential myeloid–keratinocyte signaling axis in PR. In addition, PR skin exhibited elevated expression of *IL-4* and *IL-13* alongside TWEAK signaling target genes, including *TNF, IFNG, IL-6, CCL22*, and *CCL17*, suggesting a distinct pattern of spatially coordinated immune activation in PR that is not observed in classical AD or PP. To further understand the role of myeloid cells in PR, we performed subclustering analysis of myeloid cells, revealing distinct subsets, including conventional dendritic cells (cDC1, cDC2a, cDC2b), plasmacytoid dendritic cells (pDC), Langerhans cells, and macrophages (Fig. 3G). Within these subsets, PR myeloid cells, particularly M2 macrophages and cDC2s, exhibited high *TNFSF12*, highlighting the specialized role of myeloid cells in TWEAK-mediated inflammation in PR (Fig. 3H). Comparative analysis between PR and non-PR myeloid cell transcriptomes identified numerous differentially expressed genes (DEGs), as illustrated by a volcano plot that underscored significantly upregulated mitochondrial genes, suggesting highly metabolically active myeloid cells in PR (Fig. 3I). To define the regulation of *TNFSF12* mRNA expression and its relationship to biological treatment in PP, we screened datasets from RNA-seq and microarray studies and identified that TNFα, an upstream regulator of Th17 inflammation in PP, downregulates *TNFSF12* gene expression in different types of monocytes (Figure 3J). Further, we stimulated THP-1 myeloid cells in vitro with the TNF activator lipopolysaccharide (LPS), alone or in combination with an anti-TNFα agent, infliximab. Notably, infliximab treatment significantly increased *TNFSF12* expression compared to LPS treatment alone, implicating TNF- α inhibition in enhancing TWEAK signaling (p = 0.044; Fig. 3K). These data suggest a central role for myeloid–keratinocyte interactions in PR pathogenesis, where activated myeloid cells expressing elevated *TNFSF12* directly engage their cognate receptors on keratinocytes, enhancing downstream inflammatory and remodeling responses that are characteristic of paradoxical inflammation (Fig. 3L).

**Fig 3.**
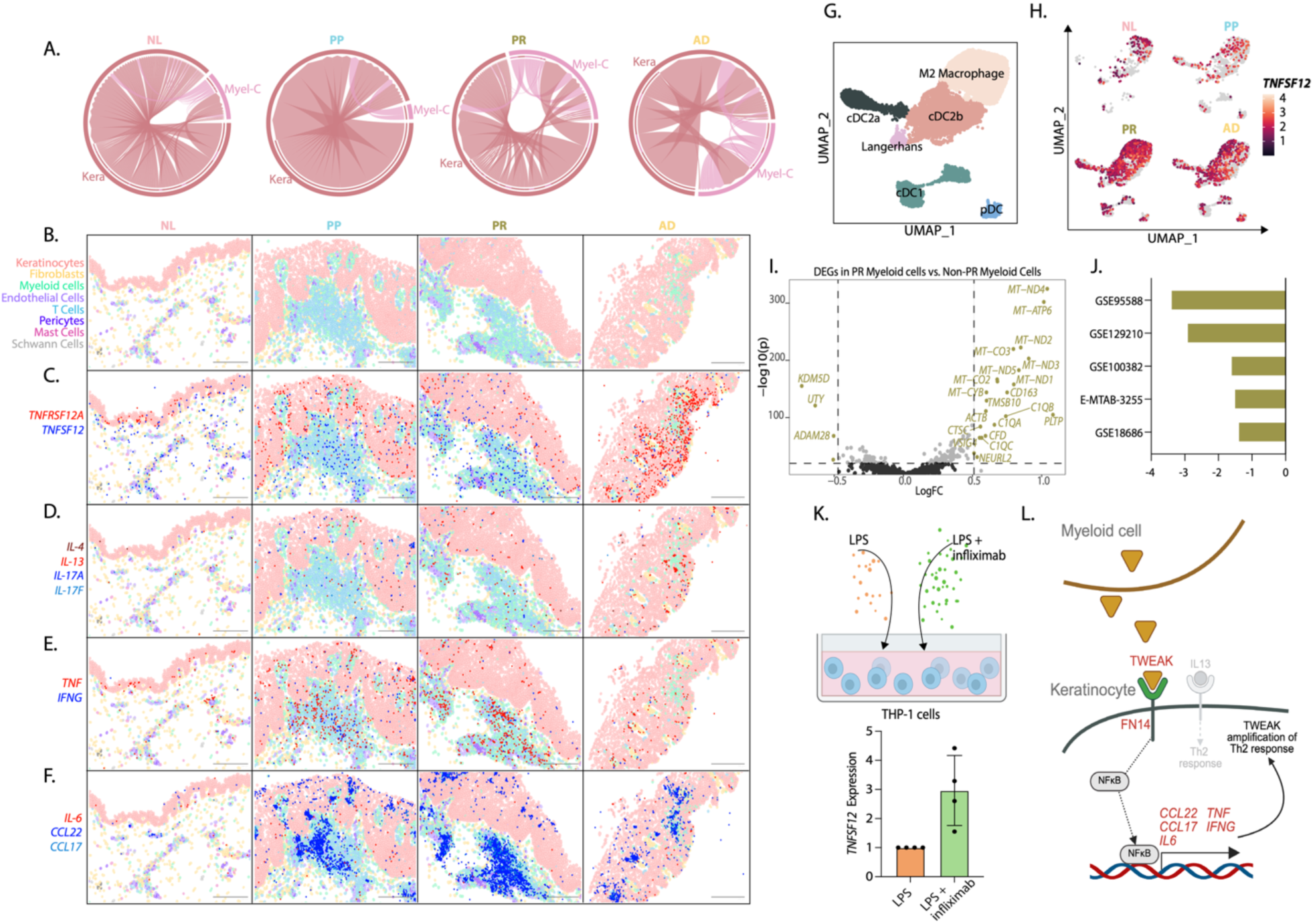
Myeloid–keratinocyte interactions drive paradoxical inflammatory responses via TWEAK signaling and enhanced cytokine communication. A. Circos plots illustrating cell– cell interactions strength specifically between myeloid cells and keratinocytes in NL, PP, PR, and AD respectively (arrow width signifies number of interactions); (B-F) Spatial transcriptomics representative images of NL, PP, PR and AD skin highlighting spatial distribution and co- localization of major cell types and gene transcripts of critical inflammatory mediators: B. Labeling of Keratinocytes, Fibroblasts, Myeloid Cells, Endothelial Cells, T Cells, Pericytes, Mast Cells, and Schwann Cells; C. *TNFSF12* and *TNFRSF12A* transcript expression; D. *IL-4, IL-13, IL- 17A,* and *IL-17F* transcript expression; E. *TNF* and *IFNG* transcript expression; F. *IL-6, CCL22*, and *CCL17* transcript expression; G. scRNA-seq UMAP of myeloid cell subclusters and annotation of distinct subsets including: conventional dendritic cells (cDC1, cDC2a, cDC2b), plasmacytoid dendritic cells (pDC), Langerhans cells, and macrophages; H. scRNA-seq feature plots across disease groups showing average expression of *TNFSF12*; I. Volcano plot comparing differentially expressed genes (DEGs) between PR-specific myeloid cells and non-PR myeloid cells; J. Bar plot depicting *TNFSF12* gene downregulation in TNFα-stimulated monocytes observed across multiple bulk-RNAseq and microarray studies (Study accession IDs reported on y-axis); K. Quantitative PCR analysis showing the normalized expression of *TNFSF12* in THP-1 monocytes stimulated with LPS alone or LPS plus infliximab; L. Working model summarizing key myeloid–keratinocyte interactions in PR skin where *TNFSF12-*expressing myeloid cells interact with *TNFRSF12A-*expressing keratinocytes, leading to amplified inflammatory signaling in PR.

### Immunohistochemical analysis validates enhanced TWEAK and FN14 expression in epidermal and inflammatory cells in PR skin

To validate and localize TWEAK signaling at the protein level in PR, we performed immunohistochemistry (IHC) for TWEAK and FN14 in NL, PP, PR, and AD skin biopsies. Across the disease groups, TWEAK staining showed a distinct expression pattern, with PR skin exhibiting increased staining intensity compared to NL and PP skin (Fig. 4A). Semi-quantitative scoring of positive and negative staining demonstrated an increased TWEAK-positive cell frequency in PR samples (Fig. 4B). Moreover, density staining analysis confirmed elevated TWEAK expression in PR and AD skin, indicating robust local upregulation of TWEAK ligand in both the epidermis and the inflammatory infiltrate (Fig. 4C). Analysis of the TWEAK receptor FN14 revealed similarly enhanced expression in PR skin, and positive versus negative staining analysis further underscored significantly higher frequencies of FN14-positive cells in PR (Fig. 4 D-E). Quantitative density staining demonstrated notably higher FN14 expression levels in epidermal keratinocytes and dermal inflammatory infiltrates in PR skin (Fig. 4F). We performed region-specific quantification of TWEAK and FN14 expression in the epidermis and dermal inflammatory infiltrates. TWEAK expression was significantly elevated in the epidermis of PR and AD lesions compared to NL and PP. Notably, FN14 expression was the highest in AD, followed by PR, with both groups showing significant upregulation relative to NL and PP, particularly within the suprabasal epidermal layers (Fig. 4G–H). Within dermal inflammatory infiltrates, PR samples similarly exhibited markedly elevated expression of both TWEAK and FN14 (Fig. 4I-J). Collectively, these immunohistochemical findings provide strong protein-level confirmation of robust TWEAK ligand–receptor expression specifically enriched in PR lesions. The pronounced epidermal and immune infiltrate expression highlights the functional role of TWEAK signaling in paradoxical inflammatory reactions, further supporting its therapeutic potential as a targeted mediator in PR.

**Fig 4.**
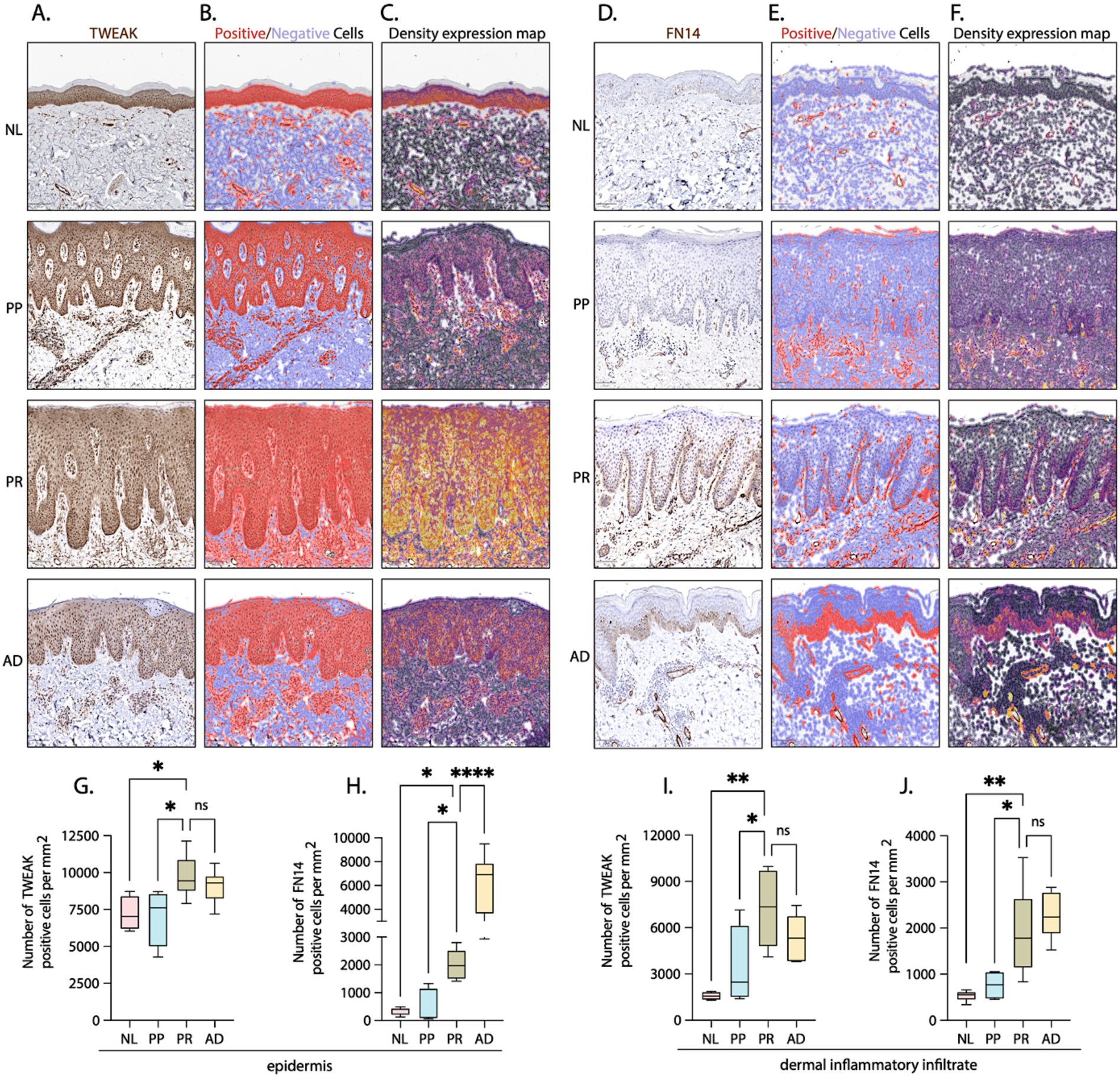
Immunohistochemical analysis validates enhanced TWEAK and FN14 expression in epidermal and inflammatory cells in PR skin. A. Representative immunohistochemistry staining images of TWEAK in skin biopsies from NL, PP, PR, and AD skin (scale bars indicate 100 µm); B. QuPath semi-quantitative staining analysis depicting TWEAK-positive (red) and TWEAK-negative (blue) cells; C. QuPath semi-quantitative staining density maps representing TWEAK staining intensity across disease groups with darker cells representing lower expression and brighter cells (reds, oranges, yellows) representing higher expression; D. Representative immunohistochemistry staining images of FN14 in skin biopsies from NL, PP, PR, and AD skin (scale bars indicate 100 µm); E. QuPath semi-quantitative staining analysis depicting FN14- positive (red) and FN14-negative (blue) cells; F. QuPath semi-quantitative staining density maps representing FN14 staining intensity across disease groups with darker cells representing lower expression and brighter cells (reds, oranges, yellows) representing higher expression; G. Quantification of TWEAK-positive cells per mm^2^ of epidermis across disease groups; H. Quantification of FN14-positive cells per mm^2^ of epidermis across disease groups; I. Quantification of TWEAK-positive cells per mm^2^ of dermal inflammatory infiltrate across disease groups; J. Quantification of FN14-positive cells per mm^2^ of dermal inflammatory infiltrate across disease groups. N = 3 skin biopsies with 2 regions of interest (ROIs) per biopsy quantified for TWEAK and FN14 staining.

### *TNFRSF12A*–expressing keratinocytes exhibit a distinct inflammatory epithelial program in PR

To investigate the epithelial response to inflammation in paradoxical reactions (PR), we performed subclustering of keratinocytes across disease groups. Unsupervised clustering revealed six distinct keratinocyte subsets: basal (Bas-KC), cycling (Cyc-KC), two differentiating (Dif-KC1, Dif-KC2), and two terminally differentiated (TD-KC1, TD-KC2) populations (Fig. 5A). We visualized the relative proportions of keratinocyte subsets and observed that Dif-KC1 was predominantly found in NL, whereas the remaining populations were more uniformly distributed across the disease groups (Fig. 5B). We next examined the expression of *TNFSF12* and *TNFRSF12A* across disease groups, specifically in keratinocytes, and observed the highest expression in AD and PR, although the proportion of keratinocytes that expressed *TNFSF12* was quite low compared to those that expressed *TNFRSF12A* (Fig. 5C). To explore the functional consequences of TWEAK receptor expression in the epidermis, we stratified keratinocytes into *TNFRSF12A*-expressing and *TNFRSF12A*-nonexpressing populations and performed differential gene expression analyses across disease groups. Among *TNFRSF12A*-expressing keratinocytes, PR displayed a distinct transcriptional program marked by the increased expression of inflammatory genes, epithelial stress mediators, and immune-interacting molecules, including cytokines, chemokines, and interferon-responsive genes (Fig. 5D). This suggests that *TNFRSF12A* expression promotes a keratinocyte phenotype uniquely poised to respond to inflammatory cues during PR. In contrast, *TNFRSF12A* non-expressing keratinocytes exhibited a separate set of differentially expressed genes across conditions, with relatively muted inflammatory signatures in PR (Fig. 5E). This opposing pattern between *TNFRSF12A*-expressing and non-expressing cells indicates that TWEAK signaling defines a functional keratinocyte state with enhanced sensitivity to inflammatory ligands and potentially a greater capacity to modulate epidermal–immune crosstalk. To further investigate the immunological state of these keratinocyte subsets, we performed Reactome “Immune System” pathway enrichment analysis in *TNFRSF12A*-expressing versus non-expressing keratinocytes under all conditions. In *TNFRSF12A*-expressing keratinocytes, the PR skin demonstrated strong enrichment of interferon signaling, including “interferon signaling,” “interferon alpha/beta signaling”, and “interferon gamma signaling” (Fig. 5F). Interestingly, *TNFRSF12A* non-expressing keratinocytes showed different immune enrichment, with overlap with PP, such as “Interleukin-23 signaling,” “Neutrophil Degranulation,” and “Interleukin-1 signaling,” and lacked PR-specific activation of these inflammatory cascades (Fig. 5G). These results suggest that *TNFRSF12A*-expressing keratinocytes could act as epithelial sensors and amplifiers of inflammation in PR, and they could act as distinct entities from TNFRSF12A non-expressing keratinocytes.

**Fig 5.**
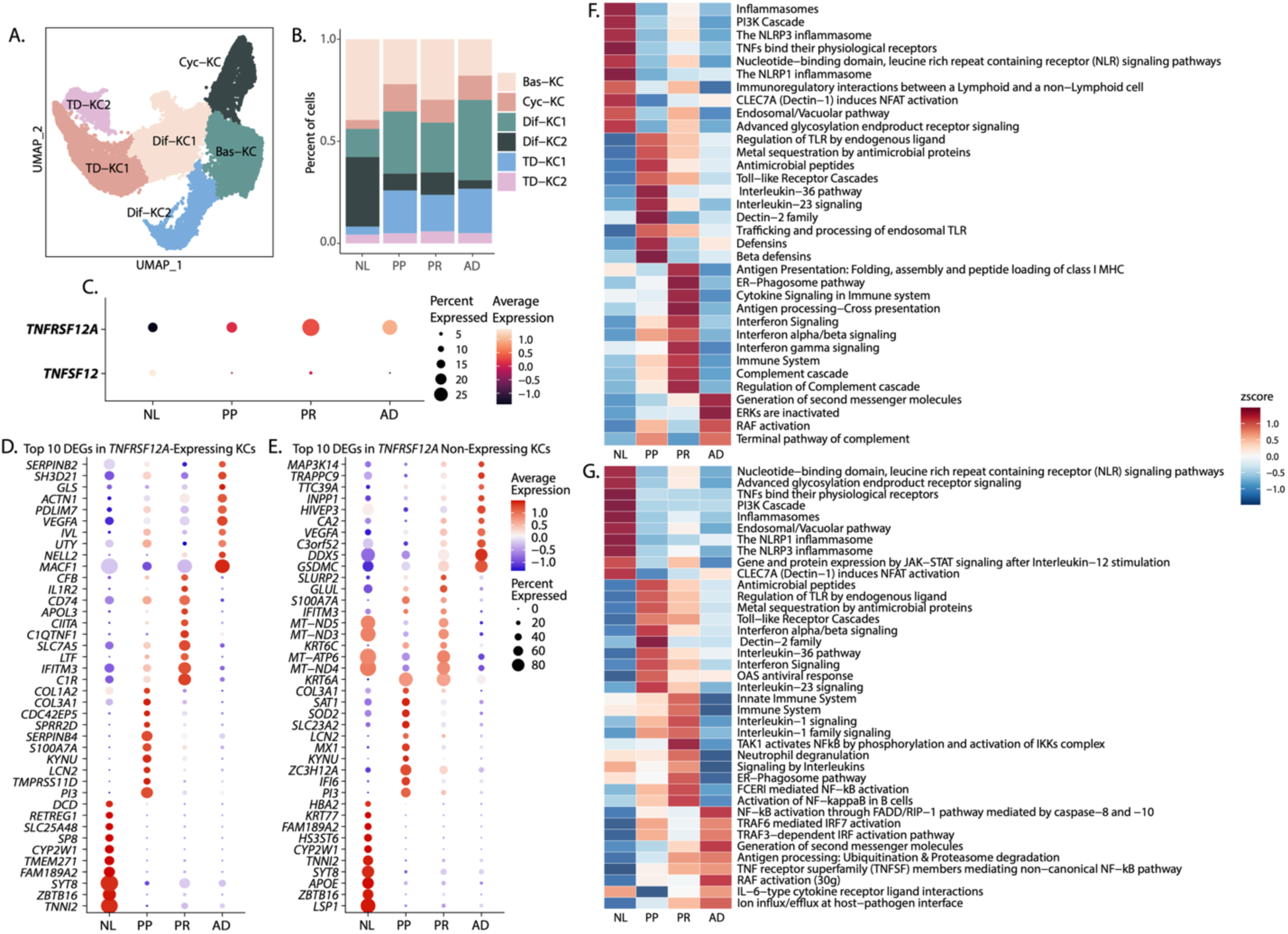
*TNFRSF12A*–expressing keratinocytes exhibit a distinct inflammatory epithelial program in PR. A. UMAP projection of keratinocyte subclusters including basal (Bas-KC), cycling (Cyc-KC), differentiating (Dif-KC1, Dif-KC2), and terminally differentiated keratinocytes (TD-KC1, TD-KC2); B. Barplot of relative proportions (percentage) of keratinocyte subsets across disease groups (NL, PP, PR, and AD); C. Dot plot showing average expression of *TNFSF12* and *TNFRSF12A* across keratinocyte subsets and disease groups (NL, PP, PR, and AD); (D-E) Dot plots of top 10 DEGs in D. *TNFRSF12A*-expressing and E. *TNFRSF12A* non-expressing keratinocytes across disease groups (NL, PP, PR, and AD); (F-G) Heatmaps representing Reactome “Immune System” pathway enrichment analysis of F. top pathways in *TNFRSF12A*- expressing keratinocytes across disease groups and G. top pathways in *TNFRSF12A* non-expressing keratinocytes across disease groups. Data is represented as a Z-score.

### TWEAK synergizes with IL-13 to activate an interferon-polarized keratinocyte program recapitulating paradoxical inflammation

To investigate the mechanistic consequences of TWEAK signaling on keratinocyte inflammatory states, we stimulated immortalized human keratinocytes (N-TERTs) with vehicle, TWEAK, IL-13, IL-17A, or a combination of IL-13+TWEAK and IL-17+TWEAK. We performed scRNA-seq to determine the transcriptional diversity of keratinocyte responses. Unsupervised clustering revealed populations corresponding to four keratinocyte states: early basal, late basal, cycling, and inflammatory/migratory phenotypes (Fig. 6A). Interestingly, the inflammatory/migratory cluster was enriched for stress response and immunomodulatory transcripts and was uniquely marked by *KRT8* and *KRT18* expression (Fig. 6B). When assessing the relative proportions of keratinocyte states across experimental conditions, we did not observe major differences, except for TWEAK and IL-13+TWEAK conditions, which notably possessed an expanded inflammatory/migratory cluster (Fig. 6C). Next, we examined differentially expressed genes across experimental groups to define the transcriptional programs associated with each cytokine condition. Compared with vehicle or single-cytokine treatment, co-stimulation with IL-13 and TWEAK induced a distinct transcriptional profile marked by elevated expression of interferon-stimulated genes (ISGs), including *ISG15*, *IFIT3*, *MX2*, and *IFIT2* (Fig. 6D). Notably, these ISGs were minimally upregulated in TWEAK stimulation alone, but not in IL-13 stimulation, highlighting a cooperative and synergistic interaction between IL-13 and TWEAK in driving a robust interferon-enriched inflammatory response. To evaluate the functional pathways downstream of this transcriptional synergy, we performed Reactome “Immune System” pathway enrichment analysis on DEGs across conditions, focusing on immune system–related pathways. The IL-13+TWEAK–stimulated keratinocytes demonstrated strong enrichment for “adaptive immune system,” “innate immune system,” “Signaling by interleukins” and importantly, “interferon signaling” (Fig. 6E). These findings suggest that TWEAK functions not only as an amplifier of Th2 cytokine signaling but also as a key switch that redirects the epithelial response toward a type I interferon–driven inflammatory state. Upstream regulator analysis further reinforced this concept by identifying IFNA2 and STAT1 as the top predicted drivers of the IL- 13+TWEAK transcriptional response (Fig. 6F). These upstream regulators are hallmarks of type I interferon signaling. This IFN-centric shift was not observed in IL-17+TWEAK–treated keratinocytes or in IL-17 alone, supporting the specificity of the TWEAK–IL-13 axis in eliciting this unique response. To assess whether this TWEAK–IL-13–induced program is relevant to paradoxical skin inflammation, we compared DEGs from IL-13+TWEAK–treated keratinocytes with DEGs of keratinocytes in PR from our patient scRNA-seq. We found 220 overlapping genes between the PR keratinocyte gene signature and IL-13+TWEAK–treated N-TERTs as well as 132 DEGs unique to IL-13+TWEAK–stimulated N-TERTs. (Fig. 6G). Pathway enrichment of both shared (n = 220 DEGs) and unique (n = 132 DEGs) gene sets revealed a strong representation of interferon signaling, cytokine–cytokine receptor interactions, and antigen processing/presentation, further validating that the TWEAK–IL-13 axis recapitulates key inflammatory features of PR keratinocytes (Fig. 6H–I). These findings suggest that TWEAK is a central amplifier of keratinocyte inflammatory response in the presence of Th2 cytokines. Through the synergistic induction of interferon-driven programs, TWEAK reprograms epithelial cell states in a manner that mirrors paradoxical inflammatory skin disease. These results offer mechanistic insights into aberrant stromal–immune crosstalk. TWEAK-expressing myeloid cells and IL-13–rich environments, can drive epithelial inflammation in PR, and suggest that TWEAK-FN14 interaction blockade may offer a therapeutic opportunity to restrain such dysregulated epithelial activation.

**Fig 6.**
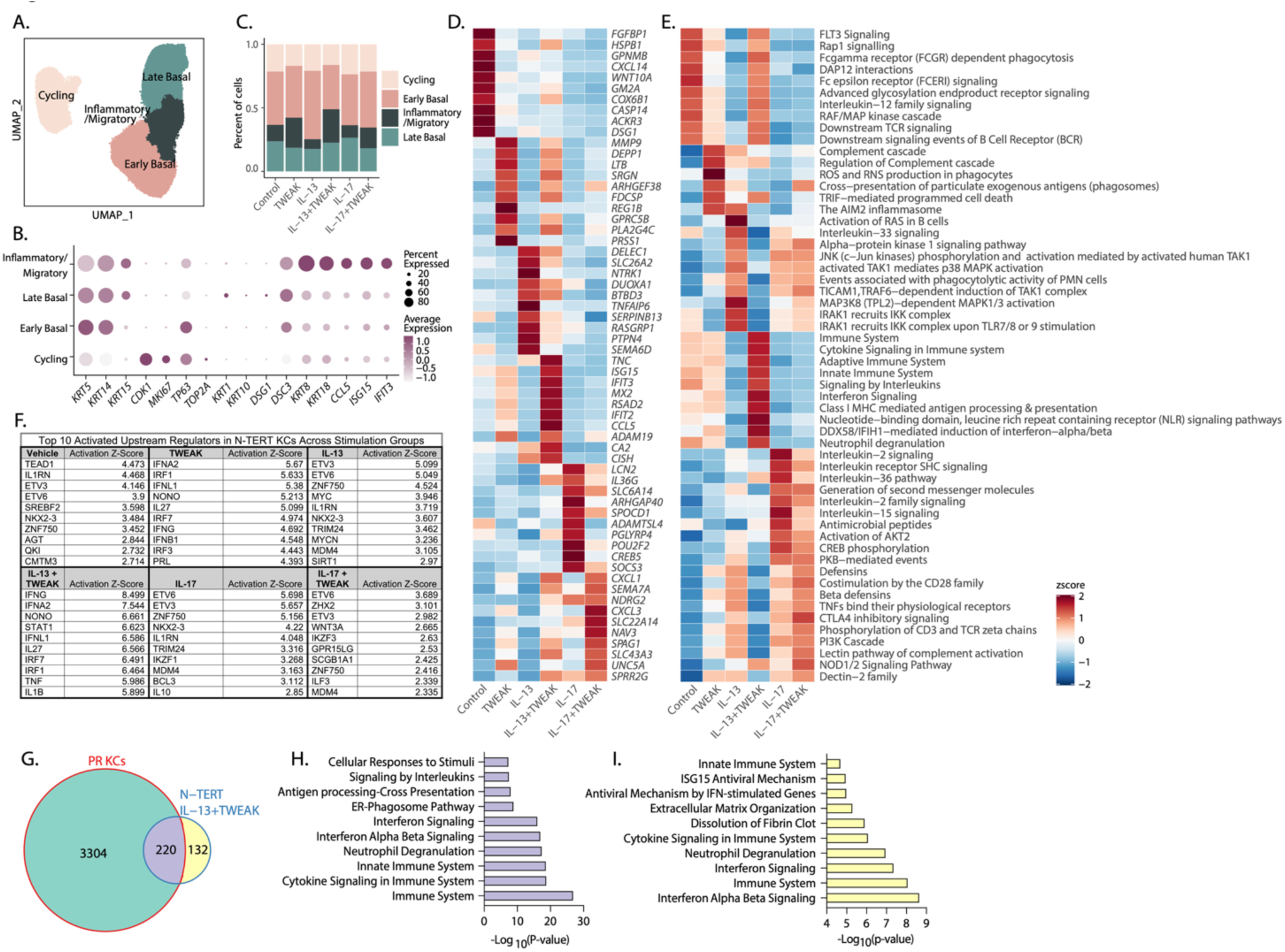
TWEAK synergizes with IL-13 to activate an interferon-polarized keratinocyte program recapitulating paradoxical inflammation. A. UMAP of N-TERT keratinocyte states including Early Basal, Late Basal, Cycling, and Inflammatory/Migratory; B. Dot plot showing average expression of keratinocyte cell state-specific gene markers; C. Bar plot showing relative proportions (percentage) of keratinocyte states across reach experimental condition; D. Heatmap of top 10 DEGs across experimental conditions; E. Heatmap of top 10 Reactome “Immune System” pathways across experimental conditions; F. Table of top 10 upregulated upstream regulators for each experimental condition; G. Venn diagram of Venn diagram showing overlap between differentially expressed genes in PR keratinocytes and IL-13 + TWEAK–treated N-TERT keratinocytes, revealing 220 shared DEGs and 132 DEGs unique to IL-13+TWEAK-treated N-TERT keratinocytes; (H–I) Reactome pathway enrichment for (G) shared and (H) IL- 13+TWEAK-treated N-TERT keratinocytes DEGs.

**Supplementary Table S1.** Paradoxical Reactions study cohort. The study cohort comprised 8 patients with paradoxical reactions (PRs). Five patients were female, and three patients were male.

| Supplementary Table S1 |  |  |  |  |  |  |  |
| --- | --- | --- | --- | --- | --- | --- | --- |
| Patients | Diagnosis | Age/Sex | Atopy | Culprit drug | Lag period (months) | Serum Eosinophils, Total IgE | Management |
| 1 | Psoriasis, Psoriatic Arthritis | 49/ F | Son w Asthma | Adalimumab | 24 | Eos-0.84<br>IgE- 632 | Golimumab + Dupilumab |
| 2 | Psoriasis, Psoriatic Arthritis | 68/ F | None | Adalimumab | 12 | N/A | Risankizumab |
| 3 | Psoriasis, Psoriatic Arthritis | 71/ F | None | Ixkezumab | 1 | Eos-1.1<br>IgE- 638 | Cyclosporine |
|  |  |  |  | Guselkumab | 1 |  | Dupilumab with arthritis flare ----- Upadecitinib |
| 4 | Psoriasis, Psoriatic Arthritis | 48/ F | Daughter w Atopic dermatitis | Ixkezumab | 2 | IgE- 112 | Switch to Risankizumab ----- Upadacitinib |
| 5 | Psoriasis | 42/ M | None | Secukinumab | 18 | N/A | Risankizumab |
| 6 | Psoriasis, Psoriatic arthritis | 76/ M | None | Secukinumab | 6 | 780 | Secukinumab +Topical corticosteroid |
| 7 | Psoriasis | 18/ F | Father w Allergic Rhinitis<br>Mother w Atopic Dermatitis | Risankizumab | 8 | IgE- 238 | Rizankizumab + Topical C.S<br>NBUVB |
| 8 | Psoriasis | 69/ M | None | Rizankizumab | 2 | N/A | Rizankizumab + Topical C.S |

Abbreviation- Topical C.S- topical corticosteroids
Normal range values for eosinophils (0.1-0.7K/□l), Total IgE (0-87 IU/ml)

Age of at PR onset ranged from 18 to 71 years (median=58.5). All patients had psoriasis vulgaris, and five of the patients also had psoriatic arthritis. Patients developed PRs while being treated with anti-TNF (2 patients), anti-IL-17 (4 patients) or anti-IL-23 (2 patients). Patient 3 had PRs during treatment with either ixekizumab or guselkumab. Lag period between starting treatment with the causative drug and PR diagnosis ranged from 1 to 24 months. Three of the patients had a personal and/or family history of atopy. Five patients had laboratory evidence of eosinophilia or elevated IgE levels. Three patients continued to be treated with the same biologic agents in addition to topical corticosteroids/phototherapy. The treatment of the others was switched to a different class.

## DISCUSSION

PR is an inflammatory skin eruption that develops in patients with chronic inflammatory diseases, such as psoriasis, during treatment with targeted biological agents, including IL-17, TNF-α, or IL- 23 inhibitors, and poses a complex clinical challenge. Using scRNA-seq, spatial transcriptomics, and in vitro stimulation assays, we uncovered a distinct immunopathogenic axis in PR driven by aberrant myeloid–keratinocyte interactions, specifically involving synergistic signaling between IL-13 and TWEAK. Our data revealed a cellular architecture in PR that was distinct from that in PP and AD. While PP is characterized by neutrophil-rich, IL-17–dominated inflammation and AD by eosinophilic, Th2-skewed responses, PR lesions demonstrate enhanced infiltration of myeloid cells in the dermis, alongside a transcriptionally reprogrammed keratinocyte compartment. These data support the hypothesis that PR constitutes a unique inflammatory entity, rather than merely a clinical overlap of PP and AD, echoing prior histologic observations of paradoxical eruptions triggered by biological treatment. Indeed, large-scale cohort studies have identified IL-17 and TNFα inhibitors as biological therapies most frequently associated with paradoxical eczematous reactions, with IL-23 inhibitors posing a lower risk (19). Additional risk factors, such as age, female sex, and prior atopic history, further underscore the need for mechanistic studies to define the trajectory of PR development (19).

Transcriptional profiling identified robust enrichment of IL-4/IL-13 signaling in PR and AD, consistent with a Th2-like inflammatory milieu. However, a defining feature of PR is selective upregulation of *TNFSF12* and *TNFRSF12A*. Single-cell mapping revealed the myeloid-specific expression of *TNFSF12* and keratinocyte-specific expression of *TNFRSF12A*. Interestingly, Keratinocytes expressing *TNFRSF12A* displayed a transcriptional profile enriched in pro-inflammatory signaling, including IFN-I signaling. *TNFRSF12A*⁺ keratinocytes appear to form a distinct epithelial subset that may serve as inflammation-amplifying cells. This concept is supported by spatial transcriptomic analyses, which showed spatial colocalization of *TNFSF12*⁺ myeloid cells with *TNFRSF12A*⁺ keratinocytes in the PR skin, reinforcing the in-situ plausibility of this signaling axis. To mechanistically validate these findings, we modeled keratinocyte responses *in vitro* using immortalized N-TERTs. Although IL-13 or TWEAK alone elicited modest transcriptional changes, their combination induced a synergistic IFN-enriched inflammatory response. Reactome “Immune System” pathway analysis and upstream regulator predictions (e.g., STAT1, IFNA2) further confirmed that TWEAK synergizes with IL-13 to reprogram epithelial cells toward an interferon-polarized state. Notably, this response was distinct from IL-17 co-stimulation, supporting the specificity of the TWEAK–IL-13 interaction. Moreover, the overlap between IL-13+TWEAK-stimulated DEGs and PR keratinocyte DEGs suggested that this axis partially recapitulates the epithelial transcriptome of PR *in vivo*. Our *in vitro* model also revealed that TNF-α blockade via infliximab potentiated *TNFSF12* expression in LPS-stimulated myeloid cells. This is consistent with findings from proteomic and single-cell analyses showing increased expression of IFN-γ, IFN-α, and TNFα in paradoxical eczema cases, along with enhanced leukocyte chemotaxis and reduced cytolytic cell activity, indicating systemic immune rewiring (20). This suggests that in the context of TNFα inhibition, compensatory upregulation of other TNFα superfamily pathways, particularly TWEAK, may perpetuate unintended inflammatory responses. These findings are consistent with the emerging literature on cytokine blockade– induced compensatory circuits, including the perpetuation of interferon responses following TNFα or IL-17 inhibition (9). Downregulation of *STAMBP* and upregulation of *IL1B* and *CCL19* in paradoxical eczema have been identified as early events following biologic therapy, highlighting systemic innate immune activation (*24*). Interestingly, gene set enrichment analyses revealed activation of Th1, Th2, Th17, and IL-10 pathways, and polygenic risk scores associated with paradoxical eczema with functional SNPs involved in TNFα and chemokine signaling rather than classic atopy-associated loci (*24*). Thus, our results provide a molecular explanation for the clinical observation of paradoxical inflammation during anti-TNF-α therapy.

Taken together, these findings suggest that the TWEAK–IL-13 axis is a central orchestrator of paradoxical skin inflammation. *TNFRSF12A*-expressing keratinocytes appear to be uniquely primed to integrate Th2 cytokine signals and myeloid-derived TWEAK, resulting in an IFN-I-dominated transcriptional state. Recent studies have further shown that IL-13 enhances TWEAK responsiveness in keratinocytes by promoting NF-κB and MAPK signaling, recruiting IFN-producing immune cells, and converging on shared downstream inflammatory mediators, such as IL-6, CXCL10, and S100 proteins (*25*). These IL-13+TWEAK co-signals also amplify the expression of disease-specific cytokines such as TSLP in AD and IL-19 in psoriasis, indicating that TWEAK could act as a disease-agnostic inflammatory amplifier (*25*). This feed-forward inflammatory loop suggests that TWEAK–FN14 signaling defines a distinct pathophysiological mechanism driving the development of PR. Importantly, the selective enrichment of TWEAK ligands and receptors in PR across transcriptomic, spatial, and *in vitro* assays suggests its potential utility as a biomarker or therapeutic target.

Despite the strengths of our multimodal approach, several limitations warrant further discussion. First, although our data highlight a compelling transcriptional and spatial correlation between TWEAK and PR, causality was not directly established *in vivo*. Although *in vitro* modeling supports functional synergy between TWEAK and IL-13, future studies using animal models or human ex vivo skin cultures are needed to confirm pathogenicity and therapeutic reversibility. Second, the exact trigger for TWEAK upregulation in patients treated with TNF-α inhibitors remains to be clarified, although our THP-1 data suggests compensatory cytokine network rewiring at the myeloid cell level. Additionally, our cohort was limited by the small sample size. Finally, while N-TERTs offer a useful model, validation in primary keratinocytes and *in vivo* tissues is necessary for translational rigor. In conclusion, our integrated study defines PR as a distinct fibroinflammatory skin disorder orchestrated by aberrant TWEAK–IL-13 signaling between myeloid cells and keratinocytes. Supported by genetic, proteomic, and single-cell data from multiple independent studies, PR has emerged as a cytokine-blockade-induced, interferon- polarized dermatosis distinct from classic Th2-driven AD. By uncovering this keratinocyte- myeloid circuit, we provide both mechanistic insights into PR pathogenesis and a foundation for the development of targeted therapies.

## MATERIALS AND METHODS

### Human samples and IRB statement

Patients and healthy controls were recruited from the Departments of Dermatology of University of Michigan and Emek Medical Center. Participants provided informed consent prior to their inclusion in the study according to a protocol approved by medical centers (IRB Protocols: EMC #0021-10, UM# HUM00174864). Patients or their legal guardians signed informed consent forms for publication of clinical photos.

### scRNA-seq of patient biopsies

The scRNA-seq dataset (106,331 cells) from patient samples was generated with 20 skin biopsies from healthy (NL), psoriasis (PP), paradoxical reaction (PR), or AD patients. Single cell RNA sequencing was performed on formalin-fixed and paraffin-embedded (FFPE) tissues using the 10X Genomics Flex-seq kit according to the manufacturer’s instructions. Tissue was dissociated per the manufacturer’s instructions using the gentleMACS Octo Dissociator. In brief, 50um scrolls were placed into a GentleMACS C-tube (Miltenyi BioTech). The scrolls were then deparaffinized, washed and dissociated into a single cell suspension enzymatically using Liberase TH. The University of Michigan Advanced Genomics Core constructed the single-cell libraries using the 10X Chromium system with chemistry. Single cell sequencing was performed on the Illumina 6000 platform to generate 150bp paired end reads.

### scRNA-seq data processing, clustering and celltype annotation

Reads were aligned to Gencode 38 (hg38) using 10X Cellranger. Quality control was performed to remove ambient RNA with SoupX v1.6.2 (*26*) and doublets with scDblFinder v1.12.0 (*27*) using default settings. Low quality cells, classified as cells with fewer than 200 genes, more than 7,500 genes, or mitochondrial gene transcripts exceeding 20%, were filtered out using the R package Seurat v4.3.0 (*28–32*). The data was normalized using the NormalizeData function and the “LogNormalize” method. The data was scaled using the ScaleData function and the top 2000 most variable genes were identified using the FindVariableFeatures function. Dimensionality reduction was performed using RunPCA on the 2000 most variable genes to derive principal components (PCs) and Uniform Manifold Approximation and Projection (UMAP) was applied to the top 30 PCs. Batch effect correction was conducted using R package Harmony v1.0 (*33*), with donor as the batch variable. Following batch correction, cells were clustered using the FindNeighbors function with shared nearest neighbor (SNN) modularity optimization-based clustering. Then the FindClusters function was performed for modularity optimization-based clustering with a resolution of 0.5 to obtain biologically meaningful clusters. For visualization, the RunUMAP function was applied to the top 30 principal components to generate a Uniform Manifold Approximation and Projection (UMAP) representation of the data. Cluster-specific marker genes were identified using the FindAllMarkers function in Seurat v4.3.0. Cell type annotations were determined by comparing these marker genes to curated cell-type signature genes.

### Differential gene expression and pathway analysis

All downstream analyses were performed in R (v4.4.1) using Seurat, ScCustomize (*34*), SeuratExtend (*35*), and CellChat (*36*) packages. We identified pseudobulk differentially expressed genes across all disease groups using the FindAllMarkers function including genes that significantly differentially expressed (adjusted p-value < 0.05), have at least ∼25% log fold-change, are expressed in at least 10% of cells in either group, and includes both upregulated and downregulated genes. We applied filters to remove low-confidence or uninformative genes, including predicted genes (e.g., AL, AC, AP) and long intergenic noncoding RNAs (LINC). We generated Venn diagrams to identify common and unique gene lists across disease groups. Genes of interest were represented as dot plots or feature plots using the DotPlot_scCustom or FeaturePlot_scCustom functions from the ScCustomize v3.0.1 package. We used the Enrichr database to identify Reactome pathways uniquely enriched in PR (*37–39*). We used the SeuratExtend package and the Reactome database to perform additional gene set enrichment analysis (GSEA), focusing on the “Immune System” parent category across disease groups. GSEA was conducted using the GeneSetAnalysisReactome function, pathway names were standardized using the RenameReactome function, and most variable immune pathways were identified using the CalcStats function, and the top enriched pathways were visualized using the Heatmap function. We performed a different pathway analysis using the Cellchat v1.6.1 R package and the rankNet function, which ranks signaling interactions based on communication strength (“weight”) across disease groups. A p-value cutoff of 0.01 was applied to identify significant disease-specific pathway enrichment, and results were visualized as stacked bar plots to highlight differences in signaling strength and interaction patterns.

### Upstream regulator analysis

DEG lists with associated log fold-changes, and p-values, were uploaded to Ingenuity Pathway Analysis (IPA, QIAGEN Inc.). Upstream Regulator Analysis was used to identify potential transcriptional regulators responsible for observed expression changes. Regulators were prioritized based on activation z-score (|z| ≥ 2) and p-value of overlap (Benjamini-Hochberg adjusted p < 0.05).

### Cell-cell communication networks

To compare intercellular communication roles across skin anatomical sites, we performed signaling role analysis using the CellChat package in R. In brief, we processed our Seurat object according to the CellChat vignette and generated CellChat objects for each disease group. We then compiled the individual CellChat objects into a list and merged using the mergeCellChat and updated the merged object using updateCellChat to harmonize metadata. To visualize intercellular signaling networks across disease groups, we used CellChat-inferred interaction probabilities and the circlize R package to generate circos plots (*40*). Cell–cell communication data was extracted using the subsetCommunication() function with a probability threshold of 0.05 from the “net” slot. Interactions were filtered for strong connections (prob > 0.05) and formatted to include sender (“from”), receiver (“to”), and interaction strength (“value”). For global signaling visualization, all significant cell–cell interactions were displayed as directional chords. To highlight specific myeloid–keratinocyte interactions, communication data were further filtered to include only communication between “Myeloid cells” and “Keratinocytes.”

### Spatial transcriptomics

Spatial gene expression analysis was performed using the 10x Genomics Xenium in situ platform with a custom-designed Xenium Human Gene Expression Panel, following the manufacturer’s protocol. FFPE tissue sections were mounted on Xenium-compatible slides and processed at the University of Michigan Advanced Genomics Core, including fixation, permeabilization, and hybridization steps. Fluorescently labeled probes targeting preselected genes were applied to enable high-resolution, single-cell spatial transcriptomic profiling. Sequential in situ hybridization and imaging cycles captured transcript localization and abundance. Raw output was processed using Xenium Ranger for image registration, cell segmentation, and transcript quantification. Data visualization and analysis were conducted using Xenium Explorer 3.

### Immunohistochemistry

Paraffin-embedded skin sections from excisional biopsies of NL, PP, PR and AD patients were baked at 60 °C for 30 minutes, deparaffinized, and rehydrated through graded alcohols (*41*). Antigen retrieval was performed in pH 9 buffer using a pressure cooker at 125 °C for 30 seconds. Following cooling, sections were treated with 3% hydrogen peroxide for 5 minutes to quench endogenous peroxidase activity, then blocked with 10% goat serum for 30 minutes at room temperature (*41*). Primary antibodies were applied overnight at 4 °C, including: TWEAK (ThermoFisher Scientific, PA5-96379; 1:100), FN14/TNFRSF12A (Abcam, ab109365; 1:100), and Goat IgG isotype control (Jackson ImmunoResearch, 005-000-003) (*41*). The next day, slides were washed, incubated with appropriate HRP-conjugated goat α rabbit secondary antibody for 30 minutes, and developed using diaminobenzidine (DAB) substrate prior to imaging (*41*).

### Immunohistochemistry quantification and analysis

IHC quantification and image analysis was performed using QuPath v0.5.1 (*42*). Regions of interest (ROIs) were manually selected, and cell detection was carried out using QuPath’s “Cell Detection” function, with optimized parameters for nucleus size, background radius, and thresholding. Cells were classified as positive (+) or negative (-) for staining based on DAB optical density using the “Positive Cell Detection” function. For each ROI, the total number of positive and negative cells, number of positive cells per mm2, and percent positivity, were determined. To assess spatial localization of signal intensity, the “Show Measurement Maps” tool was used to generate density plots across each tissue section.

### THP-1 Cell Stimulation and qPCR Analysis

THP-1 cells (TIB-2, ATCC) were seeded at a concentration of 1 × 10⁶ cells/mL in 24-well plates. And stimulated with either lipopolysaccharide (LPS; Sigma-Aldrich, Cat# L2887) 500 ng/mL, or with LPS and infliximab (Padagis, Israel) at 10 µg/mL. Following a 4-hour, cells were harvested for RNA extraction. Total RNA was isolated from the cells using the RNeasy Mini Kit (QIAGEN, Cat# 74104) in accordance with the manufacturer’s protocol. RNA concentration was measured and equalized across samples. Complementary DNA (cDNA) was synthesized from total RNA using the qScript cDNA Synthesis Kit (QuantaBio, USA). quantitative real-time PCR (qPCR) was perfrmed using gene-specific primers for TNFSF12/TWEAK (Forward: CCCTGCGCTGCCTGGAGGAA, Reverse: AGACCAGGGCCCCTCAGTGA), all purchased from Sigma-Aldrich. Reactions were carried out using SYBR Green PCR Master Mix (Applied Biosystems, Thermo Fisher Scientific) on a StepOnePlus Real-Time PCR System (Applied Biosystems, Thermo Fisher Scientific). Relative mRNA expression levels were calculated using the ΔΔCt method, with GAPDH serving as the endogenous control. Results of LPS + infliximab treated cells were expressed as fold-change relative to LPS- stimulated cells.

### N/TERT keratinocyte culture, stimulation, and scRNA-sequencing

N/TERT keratinocytes were plated at a density of 100,000 cells per well in 12-well tissue culture plates overnight and incubated at 37°C with 5% CO2. Cells were stimulated with the following cytokines for 48 hours: no-treatment control, IL-13 (10ng/mL, R&D Systems; 213-ILB-010), IL-17 (10ng/mL; R&D Systems; 213-ILB-050), TWEAK (100ng/mL; R&D Systems; 1090-TW-025), IL-13 + TWEAK in combination, and IL-17 + TWEAK in combination. After 48 hours, cells were then harvested using trypsin and resuspended in PBS for scRNA-seq. Each condition was conducted in duplicate (n=2).

## Author contributions

Conceptualization: SM, ECB, JEG

Methodology: SM, RB, RJ, PD, LAT, JEG

Investigation: SM, FJ, MH, VVD

Visualization: SM, RB, HZ, XX, ZF, RJ

Funding acquisition: ECB, JEG

Project administration: ECB, JEG

Supervision: LCT, ECB, JEG

Writing – original draft: SM

Writing – review & editing: SM, BK, PD, JMK, ACB, ECB, JEG

## Data and materials availability

Our datasets will be freely available for download through the GEO database under the accession number (to be added upon publication). All reagents used in this study are commercially available. No cell lines were generated in this study.

